# Oncogenic GNAS uses PKA-dependent and independent mechanisms to induce cell proliferation in human pancreatic ductal and acinar organoids

**DOI:** 10.1101/2023.01.16.524220

**Authors:** Ridhdhi Desai, Senthil Muthuswamy

**Affiliations:** Cancer Research Institute, Department of Medicine, Beth Israel Deaconess Medical Center, Harvard Medical School, Boston, MA 02215, USA; Islet Cell and Regenerative Biology, Joslin Diabetes Center, Harvard Medical School, Boston, MA, USA; Laboratory of Cancer Biology and Genetics, National Cancer Institute, NIH, Bethesda, MD, MA, 02215, USA

## Abstract

Ductal and acinar pancreatic organoids generated from human pluripotent stem cells (hPSCs) are promising models to study pancreatic diseases, including precursor lesions of pancreatic cancer. Genome sequencing studies have revealed that mutations in a G-protein (GNAS^R201C^) are exclusively observed in intraductal papillary mucinous neoplasms (IPMNs), one of the most common cystic pancreatic precancerous lesions. GNAS^R201C^ cooperates with oncogenic KRAS^G12V/D^ to produce IPMN lesions in mice; however, the biological mechanisms by which oncogenic GNAS affects the ductal and acinar exocrine pancreas are not understood. In this study, we use pancreatic ductal and acinar organoids generated from human embryonic stem cells to investigate mechanisms by which GNAS^R201C^ functions. As expected, GNAS^R201C^-induced cell proliferation in acinar organoids was PKA-dependent. Surprisingly, GNAS^R201C^-induced cell proliferation independent of the canonical PKA signaling in short-term and stable, long-term cultures of GNAS-expressing ductal organoids and in an immortalized ductal epithelial cell line, demonstrating that GNAS^R201C^ uses PKA-dependent and independent mechanisms to induce cell proliferation in the exocrine pancreas. Co-expression of oncogenic KRASG12V and GNAS^R201C^ induced cell proliferation in ductal and acini organoids in a PKA-independent and dependent manner, respectively. Thus, we identify cell lineage-specific roles for PKA signaling driving pre-cancerous lesions and report the development of a human pancreatic ductal organoid model system to investigate mechanisms regulating GNAS^R201C^-induced IPMNs.

## Introduction

In the pancreas, IPMNs are the most common pancreatic cystic neoplasms and are characterized as large, mucinous cystic structures that can be clinically diagnosed through abdominal imaging (Brosens et al., 2015; Sahora and Fernandez-del Castillo 2015). Genomic analysis using whole exome and next-generation sequencing of IPMN patient samples indicate that mutations in oncogenes *KRAS*^*G12V/D*^, and *GNAS*^*R201C*^ occur early during IPMN development (Brosens et al., 2015; Wu et al., 2011; Schonleben et al.,2008; Yamaguchi H. et al.,2011). Activating mutations of *KRAS* (G12D or G12V) are observed in ∼80% of cases while activating mutations of *GNAS (*codon201) are present in 66% of IPMNs respectively (Wu J. et al., 2011; Schonleben et al.,2008; Yamaguchi et al., 2011). In particular, unlike *KRAS, GNAS* mutations are exclusively observed in IPMNs, underscoring the importance of their function in IPMN development. It is not clear if the expression of GNAS mutation is sufficient to induce disease phenotypes; however, genetically-engineered mouse models (GEMMs) indicate that co-expression of GNAS^R201C^ and oncogenic KRAS^G12D^ can form cystic, IPMN-like lesions (Taki et al, 2016; Patra et al., 2017; Ideno et al, 2018). Furthermore, the use of pdx1-or ptf1a promoters drives the expression of the transgenes in embryonic pancreatic precursors, leading to genetic changes in both ductal and acinar cell lineages. Thus, how mutant GNAS functions in the acinar or ductal lineages of the exocrine compartments to regulate the initiation and progression of IPMN lesions are poorly understood.

GNAS encodes the stimulatory subunit of a heterotrimeric G-protein, Gαs, that is involved in G-protein-coupled receptor (GPCR) signaling. Under homeostatic conditions, ligand-based activation of GPCRs promotes an exchange of GDP for GTP on Gαs. GTP-bound Gαs then activates adenylate cyclase and increases cytoplasmic levels of cyclic adenosine monophosphate (cAMP) (Turan and Bastepe 2015; Weinstein et al 2004; O’Hayre et al, 2013). GNAS^R201C^ mutations maintain Gαs in an active GTP-bound state and increase cAMP levels, which upregulates protein kinase A (PKA) and other downstream signaling cascades, including cyclin nucleotide-gated ion channels and EPAC1/2 guanine-nucleotide exchange factors. Activating mutations of GNAS were first described in the context of endocrine neoplasms, particularly pituitary adenomas (Landis CA. et al, 1989; Weinstein LS. et al, 1991). Since then, they have been identified to promote aberrant growth and proliferation in small cell lung cancer and gastrointestinal cancers, including colon, gastric carcinoma, and intraductal papillary mucinous neoplasms (IPMN) (Ramms et al 2021; Wilson et al., 2010; Matthaei et al, 2011; Molin et al., 2013; Matthaei et al., 2014).

We have previously developed pancreatic ductal and acinar organoid models from human embryonic stem cell-derived pancreatic progenitor cells and used them to model pancreatic dysplasia *in vitro* and pancreatic cancer progression *in vivo* (Huang et al., 2021). We reported that oncogenic GNAS induces IPMN-associated phenotypes, including lumen dilation and Muc2 production, specifically in the ductal but not acinar organoids (Huang et al., 2021), identifying the utility of the platform to investigate lineage-specific biological mechanisms.

Here, we investigate the role played by PKA signaling in regulating oncogenic GNAS-induced effects on exocrine pancreas. GNAS^R201C^-mediated activation of PKA-signaling induced formation of Muc2-expressing cystic structures in ductal organoids, however, GNAS-induced cell proliferation was independent of the PKA signaling pathway using two independent PKA inhibitors. Interestingly, unlike ductal organoids, GNAS^R201C^ induced cell proliferation in acinar organoids was PKA-dependent, demonstrating GNAS^R201C^ uses lineage-specific mechanisms to promote proliferation in the pancreatic exocrine lineage. Co-expression of GNAS and KRAS (R201C/G12V) neither augmented nor altered PKA independence in ductal organoids. In addition, we report the establishment of long-term cultures of GNAS^R201C^-expressing pancreatic ductal organoids from pancreatic progenitors to serve as a platform for investigating mechanisms by which GNAS promotes proliferation in a PKA-independent manner. In summary, we demonstrate a lineage-specific role for PKA signaling in regulating oncogene GNAS-induced cell proliferation and demonstrate the utility of stem cell-derived organoid models for investigating cell lineage-specific mechanisms regulating pancreatic cancer initiation.

## Results

### Oncogenic GNAS^R201C^ promotes the proliferation of ductal organoids in a PKA-independent manner

We previously showed that expression of oncogenic GNAS (R201C) in HuESC-derived ductal, but not in acinar organoids, is effective in recapitulating important features of IPMN lesions in the pancreas that include lumen dilation (Huang et al. 2021, Figure 1A). To begin to understand the mechanism by which GNAS functions in these cell types, we investigated the role played by the PKA signaling pathway. Day 8 organoids were treated with H89, a PKA inhibitor, and doxycycline to induce R201C expression and were analyzed on Day 16. H89 treatment decreased both total levels of phosphorylated PKA(pPKA) substrates and phosphorylation of a specific PKA substrate vasodilator-stimulated phosphoprotein (VASP) (Figure 1B) in GNAS-expressing ductal organoids, demonstrating inhibition of PKA activity. As expected, H89 treatment reduced R201C-induced lumen expansion and Muc2 production (Figure 1C, D), confirming a role for PKA-dependent mechanism for regulating lumen expansion and mucin expression.

**Figure 1.**
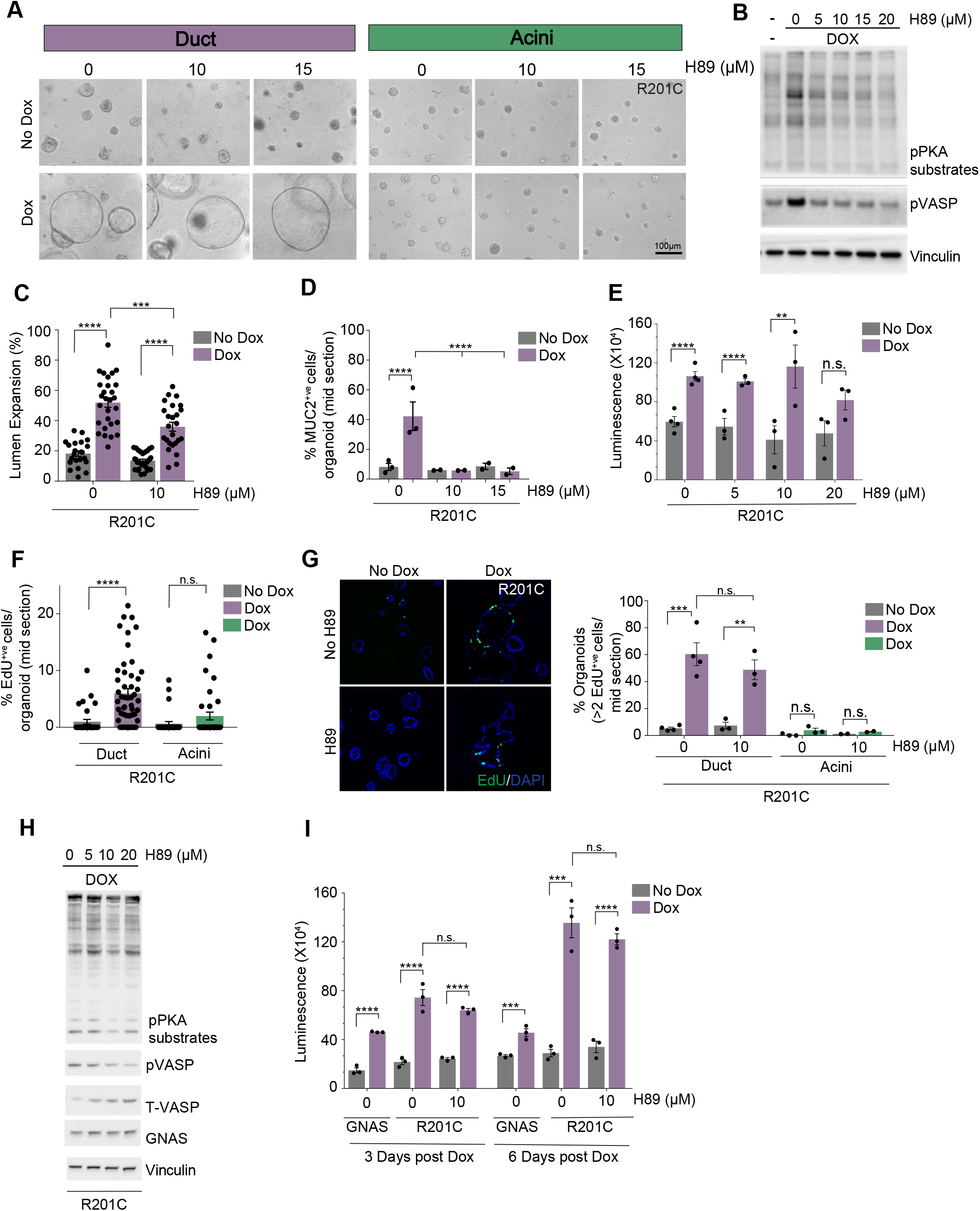
Oncogenic GNAS^R201C^ increases proliferation of ductal organoids in a PKA-independent manner (A) Phase Images (10X) of GNAS^R201C^-expressing ductal and acini organoids in the presence and absence of doxycycline with indicated doses of H89 treatment. Scale bar, 100μm (B) Western blot analysis of GNAS^R201C^-expressing ductal organoids showing inhibition of PKA activity at different doses of H89. (C) Percentage of GNAS^R201C^-expressing ductal organoids with expanded lumen in the presence and absence of doxycycline, with and without H89 treatment. (D) Percentage of GNAS^R201C^-expressing ductal organoids with MUC2 expression in the presence and absence of doxycycline, with and without H89 treatment (E) Cell titer Glow Assay measuring cell viability of GNAS^R201C^-expressing ductal organoids at different doses of H89. (F) Percentage of EdU^+ve^ cells/organoid at the midsection in GNAS^R201C^-expressing ductal and acini organoids. (G) Immunofluorescent staining of EdU^+ve^ cells in GNAS^R201C^-expressing ductal organoids, in the presence and absence of H89. Scale bar: 100μm Adjacent bar graph represents percentage of GNAS^R201C^-expressing ductal and acini organoids with at least 2 or more Edu^+ve^ cells at the midsection, with and without H89 treatment. (H) Western Blot analysis showing inhibition of PKA activity in GNAS^R201C^-expressing HPDE cells at multiple doses of H89. (I) Cell titer Glow assay measuring cell viability of GNAS^R201C^-expressing HPDE cells, with and without H89 treatment.

Activation of R201C in ductal organoids induced a ∼5.9-fold increase in the percentage of EdU^+ve^ cells compared to a ∼3-fold increase in R201C-expressing acini organoids (Figure 1F). Interestingly, we found that ∼60% of R201C-expressing ductal organoids contained two or more EdU^+ve^ cells (Figure 1G,H) both in the absence and presence of H89, identifying a PKA-independent mechanism for promoting cell proliferation. By contrast, only 3% of R201C-expressing acini organoids contained two or more EdU^+ve^ cells, which was inhibited by H89 (Figure1G,H). Oncogenic GNAS also induced a significant increase in total cell numbers, as measured by both an increase in viable cells and a total number of nuclei per ductal organoid (Figure 1F, Supplementary Figure 1A). Inhibition of PKA signaling did not significantly alter the R201C-induced increase in cell viability of ductal organoids (Figure 1F, supplementary Figure 1A).

To determine if the PKA-independent proliferation is not restricted to stem cell-derived organoids, we evaluated R201C-induced growth properties in an immortalized human pancreatic ductal epithelial (HPDE) cell line. When cultured in 3D, R201C-expression promoted an ∼1.8-fold increase in viable cells three days post Dox stimulation, and a ∼2.5 – 3.0-fold increase at six days post dox stimulation (Figure 1I). Consistent with the results obtained using stem cell-derived organoids, H89-mediated inhibition of PKA signaling did not significantly alter R201C-induced proliferative effects in HPDE cells (Figure 1H, I). Interestingly, R201C failed to induce proliferation of HPDE cells grown as monolayer cultures, suggesting a role for 3D microenvironment for R201C to promote cell proliferation of HPDE cells. Together, our findings demonstrate that oncogenic GNAS induces cell proliferation independent of PKA signaling pathway in the ductal cell lineage of exocrine pancreas.

### Concomitant expression of oncogenic GNAS and KRAS do not enhance the proliferation effects in ductal and acini organoids

In addition to GNAS mutations, mutations in KRAS are frequently found in cystic lesions of the pancreas and have been shown to co-occur with GNAS mutations in ∼50% of IPMN lesions (Wu J. et al, 2011). To address the cooperative effect between GNAS and KRAS in our stem cell model, we investigated whether oncogenic KRAS can modulate R201C-associated phenotypes in ductal and acinar structures. We generated dox-inducible lentiviral plasmids that co-express mutant forms of GNAS(R201C) and GFP-tagged KRAS(G12V) (denoted hereafter as R201C/G12V) using bicistronic expression systems (P2A peptides). Oncogenic KRAS normally localized to the cell membrane in both G12V and R201C/G12V-expressing organoids(supplementary Figure 3).

In R201C-expressing acinar organoids, expression of KRASG12V led to a substantial 19-fold increase in organoid size and a 17-fold increase in lumen expansion (Figure 2D,E,F). In comparison, acinar organoids expressing R201C alone did not induce a significant increase in organoid size but were able to expand their lumen in 20% of the organoids. KRASG12V alone produced a modest increase (∼1.9-fold) in organoid size with little to no lumen expansion in acinar organoids (Figure 2D,E,F). In ductal organoids, KRASG12V alone did not produce a significant increase in the total surface area of organoids but increased the percentage of organoids with visible lumen by ∼5-fold (Figure 2A,B,Cs). R201C-expressing ductal organoids produced a prominent increase in both organoid size (∼8-fold) and lumen expansion (5.5-fold). R201C/G12V co-expressing ductal organoids showed a significantly reduced increase in both organoid size (∼2-fold) and percentage of organoids with a visible lumen (3.4-fold) (Figure 2A,B,C).

**Figure 2.**
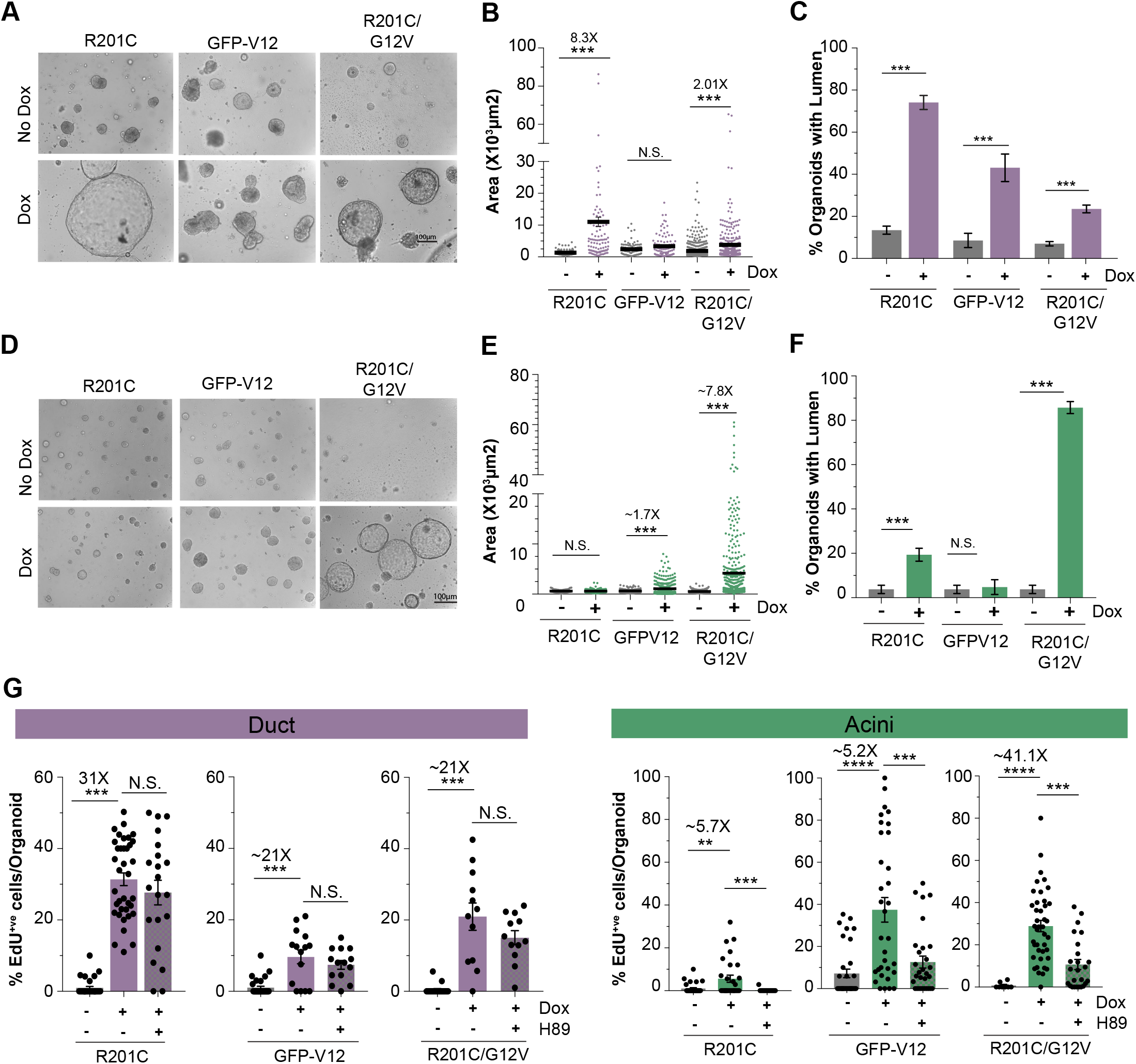
Concomitant mutations in oncogenic GNAS and KRAS do not cooperate with one another to enhance the proliferation effects of GNAS^R201C^ and KRAS^G12V^ in ductal and acini organoids. (A) Phase Images (10X) of GNAS^R201C^, KRAS^G12V^, and R201C/G12V-expressing ductal organoids in the presence and absence of doxycycline. Scale bar, 100μm (B) Scatter dot plot showing average total area of ductal organoids expressing GNAS^R201C^, KRAS^G12V^, and R201C/G12V, N=80-500 organoids, combination of three independent experiments. (C) Bar graph showing average percentage of GNAS^R201C^, KRAS^G12V^, and R201C/G12V-expressing ductal organoids with expanded lumen in the presence and absence of doxycycline, N=80-500 organoids, combination of three independent experiments. (D) Phase Images (10X) of GNAS^R201C^, KRAS^G12V^, and R201C/G12V-expressing acini organoids in the presence and absence of doxycycline. Scale bar, 100μm (E) Scatter dot plot showing average total area of acini organoids expressing GNAS^R201C^, KRAS^G12V^, and R201C/G12V, N=80-500 organoids, combination of three independent experiments. (F) Bar graph showing average percentage of GNAS^R201C^, KRAS^G12V^, and R201C/G12V-expressing acini organoids with expanded lumen in the presence and absence of doxycycline, N=80-500 organoids, combination of three independent experiments. (G) Percentage of EdU^+ve^ cells/organoid at the midsection in GNAS^R201C^, KRAS^G12V^, and R201C/G12V-expressing ductal and acini organoids, with and without H89 treatment.

Co-expression of R201C/G12V increased the proliferation of ductal structures, as measured using EdU incorporation, albeit at a slightly lower rate than R201C-expressing ductal organoids (Figure 2G). Interestingly, inhibition of PKA signaling using a PKA inhibitor (H89) did not attenuate the proliferation effects induced by R201C/G12V, indicating that oncogenic GNAS, either alone or in combination with KRASG12V promotes cell proliferation of ductal epithelia through PKA-independent mechanisms (Figure 2G). In acini organoids, we observed a significant ∼8.5-fold and 6.6-fold increase in both KRASG12V and R201C/G12V-expressing acini structures compared to organoids expressing R201C alone (Figure 2G). Interestingly, in contrast to our observations in ductal organoids, inhibition of PKA signaling decreased the proliferation of acini organoids that express either R201C or KRASG12V or both R201C and KRASG12V, indicating that oncogenic GNAS, either alone or in combination with KRASG12V promotes cell proliferation in a PKA-dependent manner in the acinar lineage (Figure 2G).

### Establishment of long-term cultures of GNAS^R201C^ - expressing ductal organoids

We next sought to establish an R201C expressing ductal organoid stable culture model for pursuing mechanistic studies. While control ductal organoids did not expand beyond the first passage, R201C-expressing ductal organoids could be expanded for at least 15 passages (∼5-6 months). Western blot analysis showed that R201C-expressing ductal organoids at later passages (P6) promoted an increase in both pPKA and pVASP, similar to that observed at the initial passage (P0), indicating that stable cultures of R201C ductal organoids maintained its ability to activate PKA signaling (Figure 3A). Further, stable R201C-expressing ductal organoids maintained their ability to induce lumen expansion and Muc2 expression (Figure 3A,B, supplementary Figure 4).

**Figure 3.**
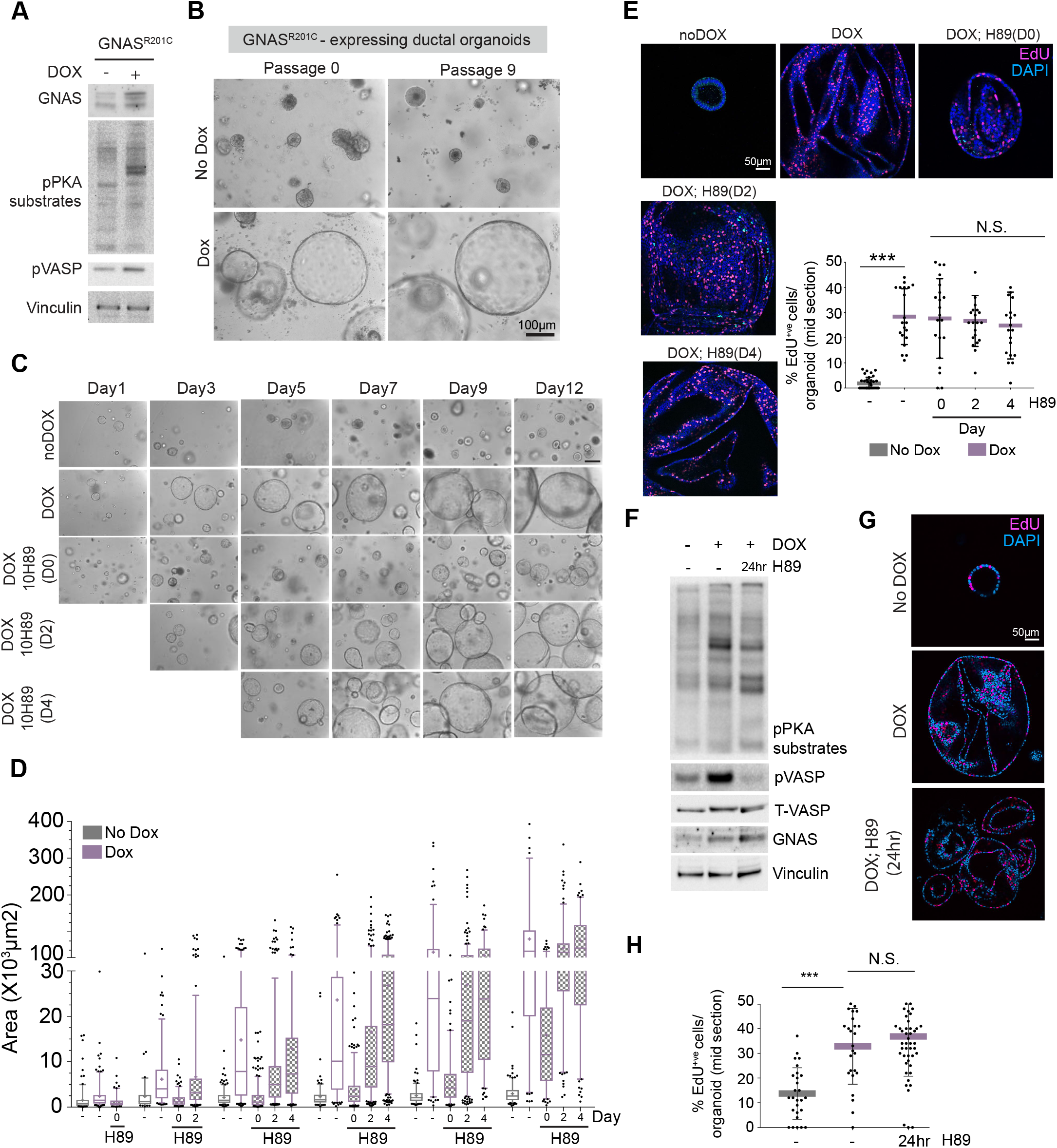
Characterization of stable R201C expressing ductal organoids (A) Western blot showing activation of PKA signaling in stable R201C expressing ductal organoids. (B) Phase images of R201C-expressing ductal organoids showing lumen expansion at multiple passages. (C) Phase images of R201C ductal organoids treated with H89 (10μM) on Day 0 (D0), Day2 (D2), Day4 (D4). Doxycycline was used to express R201C from D0. Scale Bar: 100μm (D) Box and whiskers plot showing changes in the surface area of R201C-expressing ductal organoids treated with H89 on D0, D2, and D4. (E) Immunofluorescent images of R201C-expressing ductal organoids showing proliferative organoids labeled with EdU (red) in the presence and absence of dox (from D0), when treated with H89 from D0, D2. And D4. Scale bar, 50μm. Scatter plot shows average percentage of Edu^+ve^ cells/organoid at the mid section. Lines and error bar represent mean±S.D, n=3 independent experiments. (F) Western blot showing inhibition of PKA activity after 24hr of H89 treatment. (G) Immunofluorescent images of R201C-expressing ductal organoids showing proliferative organoids labelled with EdU (red) in the presence and absence of dox (from D0), when treated with H89 for 24hrs. Scale bar, 50μm. (H) Scatter plot shows the average percentage of Edu^+ve^ cells/organoid at the mid section. Lines and error bar represent mean±S.D, n=3 independent experiments.

Next, we tested whether the growth of stable R201C-expressing ductal organoids was PKA-independent. We treated R201C-expressing ductal organoids with PKA inhibitor H89(10uM) at different stages of organoid growth (Figure 3C). We found that treatment of ductal organoids during the first 2 days (D0 or D2) of seeding briefly attenuated the growth of ductal organoids but maintained their ability to expand at 5.3-fold and 18.9-fold, respectively (Figure 3D). Whereas treatment with a PKA inhibitor on Day 4 did not have any impact and the organoid growth was comparable to untreated R201C-expressing organoids (Figure 3D). Furthermore, treatment of H89 on Day0, 2, or 4 did not affect the average percentage of Edu+ cells within each organoid (Figure 3E), suggesting that the PKA pathway is not required for oncogenic GNAS-induced proliferation both at early and late stages of organoid growth. To rule out long-term effects of H89 treatment, we monitored the effects of within the first 24 hrs of treatment. H89 inhibited PKA signaling within 24 hrs, as monitored by the reduction of p-PKA substrates and pVASP; however, this treatment did not significantly affect the percentage of EdU+ cells/organoid (Figure 3F,G, H). Lastly, to rule out inhibitor-specific effects, we treated R201C-expressing organoids with KT-5720, another PKA inhibitor, and observed no significant effect on the percentage of Edu+ structures (supplementary Figure 3). Thus, we have established stable cultures of R201C-expressing human ductal organoids, which retain both PKA-dependent IPMN-associated phenotypes and PKA-independent mechanisms to induce cell proliferation.

## Discussion

Using lineage-committed organoids, we report a surprising finding that targeted expression of mutant GNAS^R201C^ can promote the proliferation of ductal organoids through mechanisms independent of PKA signaling. By contrast, canonical Gαs-cAMP-PKA signaling is required for GNAS^R201C^-dependent proliferation of acini organoids. Our study also demonstrates that concomitant expression of KRAS^G12V^ and GNAS^R201C^ does not enhance GNAS^R201C^-induced proliferation in ductal or acinar organoids. Finally, we report the feasibility of growing GNAS^R201C^-expressing ductal organoids as stable cultures for at least 15-18 passages. These stable GNAS^R201C^-expressing ductal-like organoid cultures are an ideal platform to elucidate how GNAS^R201C^ induces cell proliferation in a PKA-independent manner and identify regulators of IPMN progression.

The kinases and downstream signals that regulate oncogenic GNAS-induced proliferation are only partly understood. Although H89 is known to inhibit activities of AKT, RSK, AMPK, and ROCK (Limbutara K. et al, 2019), it does not impact our conclusion that GNAS promotes cell proliferation independent of PKA activity because H89 efficiently inhibits PKA activity. Although cAMP-mediated activation of PKA-signaling is the major pathway downstream of GNAS activation, evidence from previous reports suggests that activation of PKA does not fully mimic the action of cAMP and, therefore, other PKA-independent pathways exist that allow cAMP to stimulate proliferation (Dremier et al, 2000; Cass et al., 1999; Dremier 1997). In addition to the activation of PKA, cAMP can activate additional downstream effectors that include EPACs, and cyclic nucleotide-gated ion channels (Cheng X et al, 2008; de Rooij J et al., 1998; Kawasaki H. et al., 1998). In this regard, our experiments indicate that treatment of organoids with Forskolin, which activates most downstream effectors of cAMP, produces more EdU+ cells than treatment with cAMP analogs that specifically activate either PKA signaling or EPAC (supplementary Figure 1). Additionally, PKA mutations are not commonly observed in gastrointestinal cancers, suggesting that at least in tissues where GNAS mutations occur, GNAS may activate additional targets in addition to PKA. Understanding downstream signals that regulate PKA-independent signaling will be the subject of further investigation.

To establish ductal and acini organoids as a disease model, we asked whether common IPMN driver mutations, KRAS and GNAS, can induce a phenotype within these cell types. Although concomitant mutations in KRAS/GNAS do not appear to enhance mutant GNAS and mutant KRAS-induced proliferation of ductal and acini organoids in culture, R201C/G12V double mutants exhibit morphological changes that are distinct from individual mutations in both ductal and acini organoids and hence provide a new platform for identifying alterations that cooperate to promote tumor progression. Our findings validate the utility of our organoid platform to gain new insights into phenotypes associated with pre-cancerous lesions in the pancreas, and future studies aimed at identifying how additional genetic alterations cooperate with R201C/G12V-to induce cancer progression in culture or in vivo will provide critical insights for the initiation and progression of PDAC and serve as a platform for identifying biomarkers of cancer progression.

## Materials and Methods

### Cell lines and Cell culture

#### Embryonic Pluripotent stem cells

In this study, HUESC8 stem cells were differentiated towards pancreatic progenitors stage in the lab of Dr. Douglas Melton, Harvard University using previously described protocols (Huang et al., 2021, Leite et al, 2020, Pagliuca et al., 2014).

#### HPDE cells

The previously described HPDE cell line was obtained from Dr. Ming Tsou (University of Toronto). HPDE cells were maintained in DMEM-HG (Gibco) supplemented with inactivated FBS and antibiotics at 37°C in 5% C0_2_. To culture HPDE cells in 3D, 8-well falcon chamber slides were coated with ∼15μl of 100% Matrigel (Growth factor reduced, Corning) and incubated at 37°C for 30mins to solidify. ∼40,000 cells were seeded in each coated well in media containing 2% Matrigel. On Day4, 1μg/ml doxycycline, with or without H89 (10μM or otherwise as indicated) was added to induce gene expression and cell viability was measured using cell titer glow 3D using manufacturer’s instructions (Promega) on Day7 and Day10. Culture media was replaced with fresh media on Day7 with doxycycline.

### Induction and culture of ductal and acini organoids

Detailed description of ductal and acini organoid induction from pancreatic progenitors is previously described (Huang L et al., 2021). Cell culture plates or 8-well falcon chamber slides were coated with undiluted Matrigel and incubate at 37°C to solidify. Transduced or uninfected pancreatic progenitors were seeded as single cells on Matrigel-coated plates in appropriate differentiation media (described in Huang et al., 2021) containing 5% Matrigel. Media change was performed every 4 days.

For long-term culture of GNAS^R201C^-expressing organoids, Day16 ductal organoids were split in a 1:3 ratio using 1mg/ml Collagenase/Dispase (Sigma) in DMEM-HG (Gibco) for 30min-1hr, followed by mechanical dissociation of organoids using a p1000 pipette. After centrifugation (1000rpm, 5min), dissociated organoids were embedded in 100% Matrigel and seeded as 80μl domes in 24-well cell culture plates. The domes were allowed to solidify for 30min at 37°C. Stage 4 ductal differentiation media containing doxycycline (1μg/ml) was used to culture GNAS^R201C^-expressing organoids. Culture media was replaced every four days and organoids were split in a 1:3 ratio every 12-16days.

### Chemicals, Drugs, and Antibodies

All reagents used for ductal and acini organoids are previously described (Huang et al., 2021). Forskolin (1099), KT-5720 (1288), and 8-CpT-0-Me-cAMP (1645) were from Tocris. H89 (S1582) was from Selleckchem; 8-Br-cAMP (B7880) was from Sigma.

The following antibodies were used: GNAS (371732, Sigma), GFP (632381,JL-8, Takara), p-VASP (3111, cell signaling), T-VASP (3132, cell signaling), pPKA substrate (9624, cell signaling), Vinculin (13901, cell signaling)

### Plasmids and Cloning

All oncogenes were cloned into pInducer21 (addgene). GNAS^R201C^ was generated as previously described (Huang et al,, 2021)

To generate pINGFPKRASG12V vector, GFPKRASG12V was amplified by PCR from pQCXIPGFPKRASG12V (Magudia et al, 2012) using primers that contain Age1 and Not1 at the N-and C-terminus: Forward, 5’-ACCGGTCGCCACCATGGTGAGCAAGGGCGAGG-3’; Reverse, 5’-GCGGCCGC TTACATAATTAC-3’, the PCR products were then subcloned into pINducer21 using Age1 and Not1.

To generate pINR201C/G12V vector, GNAS^R201C^ was amplified from using a forward primer containing Age1 and a reverse primer containing P2A sequence followed by a Not1 restriction site: Forward, 5’-5’ACC GGT CGC CAC CAT GGG3’; Reverse, 5’-GCGGCCGCCTCCGGGACCGGGGTTTTCTTCCACGTCTCCTGCTTGCTTTAACAGAGAGAAGTTCGTGGCTCCGGAGCCGAGCAGCTCGTACTGACG3’. Both PCR products were then subcloned into pINducer21 using Age1 and Not1. All final plasmids were sequence verified.

### Virus production and infection

Lentiviruses were produced in 293T cells using packaging plasmids (pCMV-dR8.9 and pMD2.G). Viral particles were concentrated using Lentivirus precipitation solution (ALSTEM). Viral pellets were resuspended in PBS and frozen as aliquots at -80°C.

To infect HPDE cells, 150,000 cells were seeded per well of a 6-well plate. Concentrated lentiviral particles were added to the cells and spinfected at 2250rpm for 30min. The viral media was then discarded. The cells were washed once with PBS and fresh culture media was added. Cells expressing the transgene were selected 48hrs after infections with 2μg/ml puromycin.

To infect pancreatic progenitors, 48-well plates were first coated with 5% Matrigel for ∼2-3hours. At the time of seeding, the Matrigel solution was aspirated and the plates were allowed to air dry. ∼150,000 single pancreatic progenitor cells were then seeded as a monolayer in each Matrigel-coated well. After 24hours, lentiviruses supplemented with 4μg/ml polybrene was added to each well and the plates were incubated at 37°C overnight. The next day, infected cells were briefly digested with Accutase (Sigma) and centrifuged at 1500 rpm for 5min. The pellet was resuspended in stage 1 of either ductal or acini differentiation media and ∼40,000 cells were seeded per well of an 8-well chamber slide. 0.3μg/ml of puromycin was added on Day3 post seeding to select for transgene-expressing cells. 1μg/ml of doxycycline was added to the culture media to induce transgene expression at indicated time points.

### Phase contrast image analysis

For phase contrast imaging, at least 50-100 organoid images (per experiment) were taken using either a 4X or a 10X objective. Image Analysis were performed using ImageJ. Briefly the longest diameter (d) of each organoid was measured and the area was calculated as π (d/2)^2^.

### Click-it EdU proliferation assay and Immunofluorescence

For Edu proliferation assays, ductal and acini organoids were incubated with 10μM Edu for 4hrs at 37°C in the incubator. Both ductal and acini structures were fixed in 4%PFA in PBS for 30min at RT and then processed for Edu-Click-iT according to manufacuturer’s instructions (Thermo Fisher). Next, the organoids were then processed for GFP immunofluorescence. Briefly, organoids were incubated with immunofluorescence buffer (7 mM sodium dibasic heptahydrate, 3 mM sodium monobasic monohydrate, 131 mM NaCl, 0.1% BSA, 0.02% Tween 20, and 0.2% Triton X-100) for 15 min at RT. Cells were then incubated in blocking buffer containing 0.5% Triton X-100 and 1% BSA for 15 min at RT. Primary antibody incubation was prepared in PBS with 1% BSA and incubated overnight at 4°C. Cells were then rinsed with PBS 3× for 10 min before secondary fluorescent antibody incubation in PBS with 1% BSA along with DAPI stain to mark the nuclei. Embedded structures were mounted on coverslips using fluorescent mounting medium (Dako). Confocal images of the mid-section of each organoid were taken on a Zeiss LSM880 inverted live-cell microscope. Images were processed using Zeiss Zen and ImageJ was used to determine percentage of Edu^+ve^ cells at the mid-section of each organoid.

### Protein Analysis and Western Blotting

For western blotting, ductal and acini cells were collected in 15ml falcon tubes by resuspension in ice-cold PBS. The suspension was centrifuged at 3500rpm for 5min. The resulting pellet was then resuspended in ice-cold cell recovery media and incubated for 30mins at 4C. The tubes were then centrifuged at 3500rpm for 5mins. The supernatant was discarded and the pellet was washed with ice cold PBS followed by centrifugation at 3500rpm for 5min. The resulting pellets were lysed in RIPA buffer supplemented with phosphatase and protease inhibitors (Roche). Protein concentration was determined by Bradford assay and standard procedures were used for western blotting. Primary antibodies used for western analysis: pPKA substrates (1:1000), pVASP (1:500), T-VASP (1:500), Vinculin (1:10000), GNAS (1:1000), GAPDH (1:2000).

### Statistics and Reproducibility

For most experiments, data was collected from at least 3 independent (i.e. three independently transduced differentiations) experiments. All quantitative data are expressed as mean ± s.e.m, unless indicated otherwise. Differences between groups were assayed by two-tailed student t-test using Prism 5 (GraphPad software). All quantitative data were collected from experiments performed in at least three samples or biological replicates. Significant differences are defined as follows: * P<0.05, **P<0.01, ***P<0.001, ****P<0.0001.

---

**Supplementary Figure 1.**
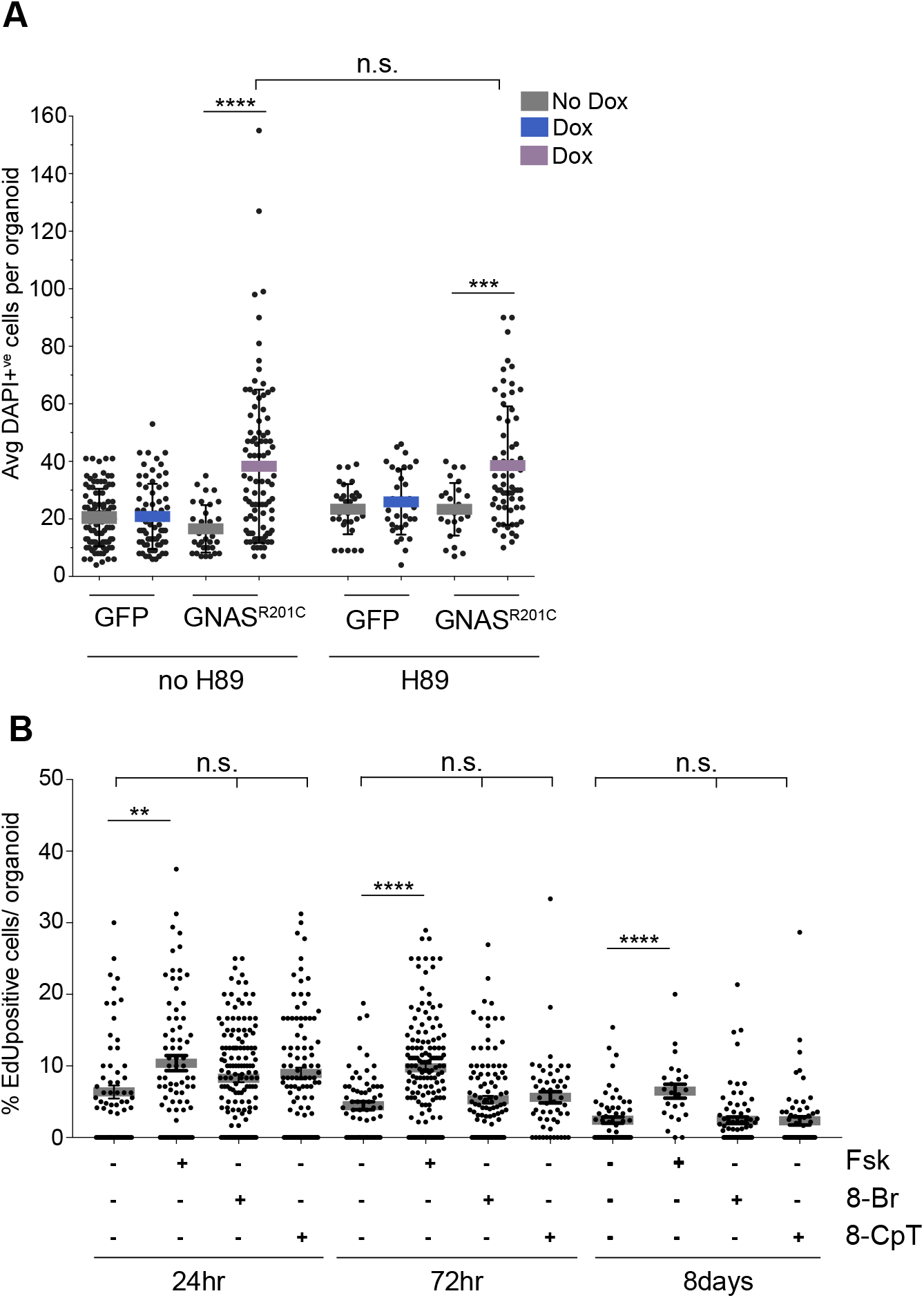
(A) Scatter dot plot showing average number of DAPI^+ve^ cells per organoid in R201C-expressing ductal organoids treated with H89(10μM), N=30-100 organoids, combination of three independent experiments. Error bar, mean±S.D (B) Scatter dot plot showing percentage of Edu^+ve^ cells per organoid in day 8 ductal organoids treated with Forskolin (10μM), 8-Br-cAMP (100μM), and 8-CpT-0-Me-cAMP (μM) for 24hrs, 72hrs, and 8days. Error bar, mean±S.D

**Supplementary Figure 2.**
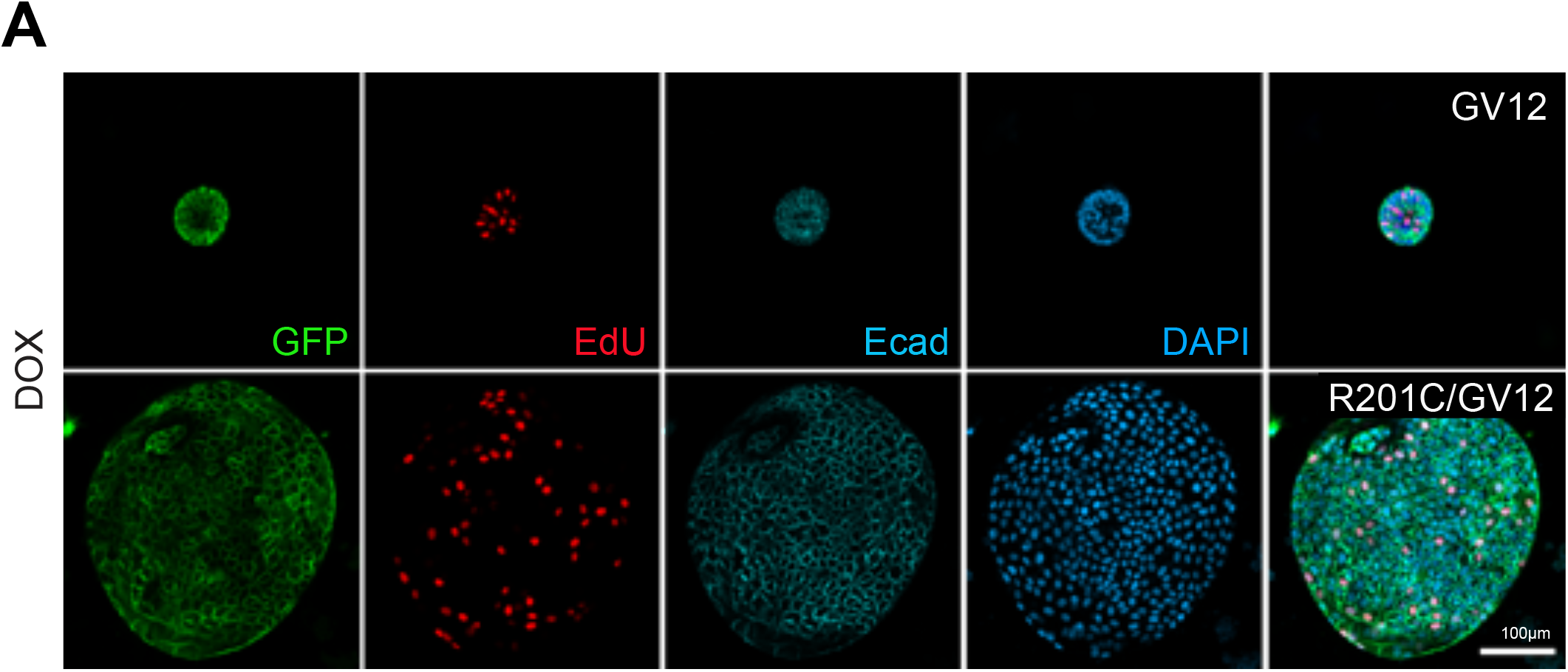
(A) Immunofluorescent images of GV12 and RC/G12V-expressing acini organoids showing membrane localization of KRASG12V. KRAS protein is marked by GFP that is tagged at the N-terminus of KRASG12V. proliferating cells are marked with EdU (red), Ecadherin (membrane marker), and DAPI (nuclei).

**Supplementary Figure 3.**
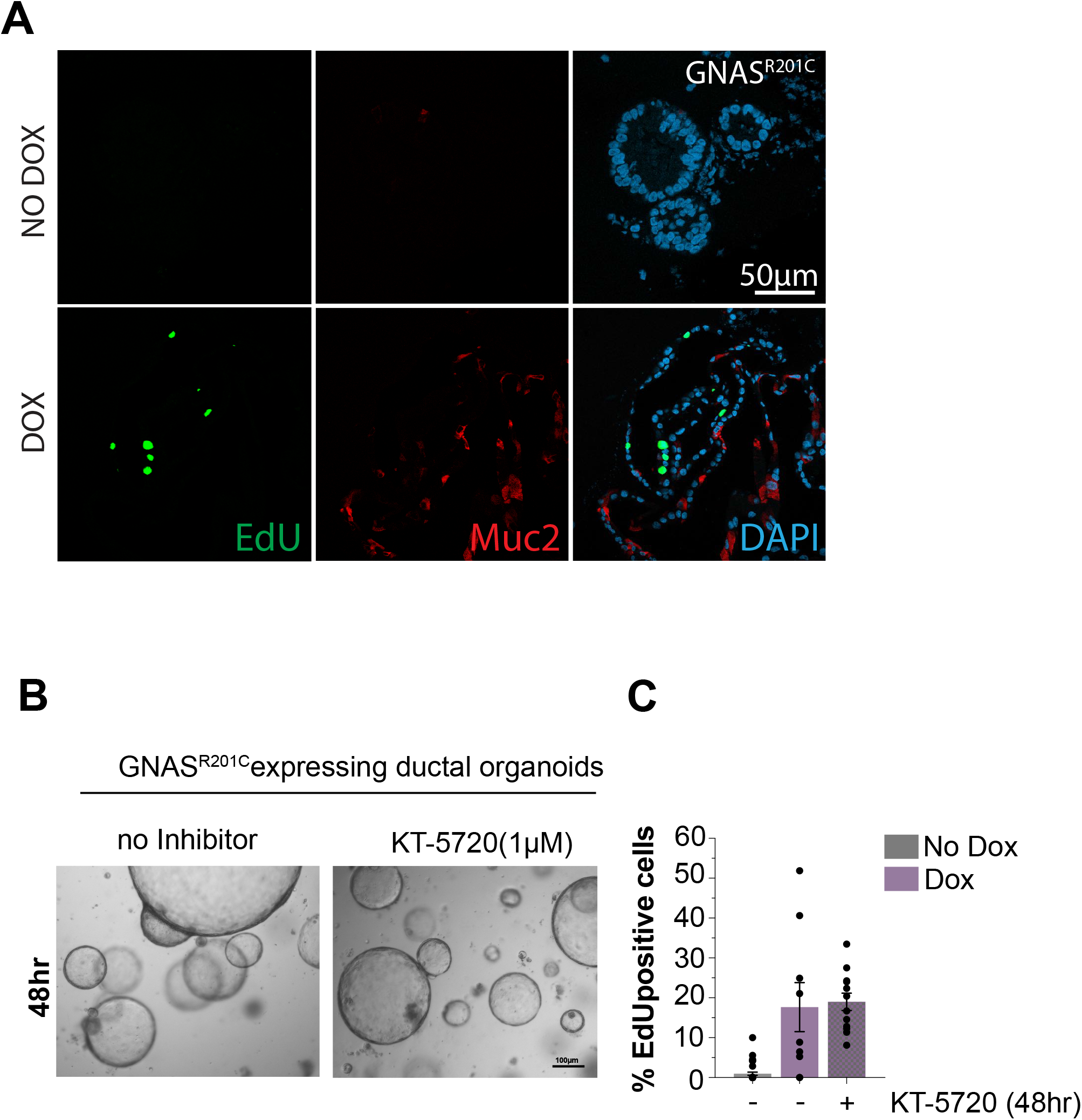
(A) Immunofluorescent staining of stable R201C-expressing ductal organoids showing proliferative cells (EdU, green), and expression of MUC2 (red), scale bar 50μm (B) Phase Images of R201C-expressing ductal organoids treated with KT-5720 (1μM) for 48hrs. (C) Bar graph showing percentage of Edu+ve cells / organoid in R201C-expressing ductal organoids treated with KT-5720 (1μM) for 48hrs.

